# Multi-omics analyses reveal rumen microbes and secondary metabolites that are unique to livestock species

**DOI:** 10.1101/2023.08.21.554168

**Authors:** Victor O. Omondi, Geoffrey O. Bosire, John M. Onyari, Caleb Kibet, Samuel Mwasya, Vanessa N. Onyonyi, Merid N. Getahun

## Abstract

Ruminant livestock like cattle, sheep, goat, and camel, have a unique digestive system with complex microbiota communities that facilitate feed conversion and production of various secondary metabolites including greenhouse gases, which are significant in livestock-vector and livestock environment interactions. However, there is limited understanding of the diversity of rumen microbes and secondary metabolites that have advantageous traits to livestock physiology, productivity, climate, and defense across different ruminant species. In this study using metagenomics and metabolomics data from four evolutionary distinct livestock species, we show that there are signature microbes and secondary metabolites for each species. For instance, camels host a unique anaerobic fungus(F) called *Oontomyces*, cattle harbor more unique microbes like *Psychrobacter* (F) and three unique bacteria genera *Anaeromyces*, *Cyllamyces*, and *Orpinomyces*. Goats have *Cleistothelebolus* (F), while sheep host *Liebetanzomyces* (F). This phenomenon may indicate that there are species-specific microbes that requires host rumen-microbes’ environment balance. Additionally, there are conserved core bacterial microbes present and in equal abundance regardless of the host genetics, indicating their essential role in maintaining crucial functions. The studied livestock fed on diverse plant materials, including grass, shrubs to acacia trees. Regarding secondary metabolites camel rumen is rich in organic acids, goat with alcohols, and hydrocarbons, sheep with indoles and cattle with sesquiterpenes. These results have implications for manipulating the rumen environment to target specific microbes and secondary metabolite networks, thereby enhancing livestock productivity, resilience, reducing susceptibility to vectors, and environmentally preferred livestock husbandry.

**IMPORTANCE:** Rumen fermentation that depends on feed component and rumen microbes plays a crucial role in feed conversion and production of various metabolites, important for physiological functions, health and environmental smartness of ruminant livestock, in addition to providing food for humans. However, given the complexity and variation of the rumen ecosystem and feed of these various livestock species combined with inter-individual differences between gut microbial communities, how they influence the rumen secondary metabolites remains elusive.

Using metagenomics and metabolomics approaches, we show that each livestock species has signature microbe(s) and secondary metabolites. These findings may contribute towards understanding rumen ecosystem, microbiome and metabolite networks, that mayprovide a gateway to manipulate rumen ecosystem pathways towards making livestock production, efficient, sustainable and environmentally friendly.

## INTRODUCTION

Livestock are an important part of the ecosystem, especially they are a major driver in most rural landscapes, diversifying belowground microbes, soil health, function, fertility and crop productivity. Globally more than 1.2 billion people are making a living in the livestock sector across the various value chains (1, 2). Ruminant livestock provide humans with foods, such as milk and meat from non-human-edible plant material, even in arid and semi-arid ecologies, where crop production is not possible due to erratic rain fall and frequent drought, thus the only means to sustainably use such vast land is through sustainable livestock husbandry. The rumen, a large fermentation chamber in ruminant livestock, harbors diverse and complex microbial communities that play crucial roles in the digestion and fermentation of feedstuff (3, 4) and production of diverse metabolites including greenhouse gases (5, 6, 7, 8). How ever, livestock vary in their resilience, and feed conversion efficiency. For instance, one-humped camel (*Camelus dromedarius*), is the most efficient and resilient animal well adapted under scarce resources in arid and semi-arid ecologies, this is recently evidenced as pastoralists shifted from cattle to camel keeping even at higher altitudes (9, 10, 11, 12). This can be taken as a climate change adaptation strategy and has potential to improve livestock climate resilience if the underlining mechanism is understood. However, the underlying mechanisms responsible for the observed variations in resilience between different livestock is not clear. We hypothesize livestock vary in their rumen microbes and secondary metabolites that has useful traits for livestock resilience and efficiency. As rumen environment hosts the most complex diverse microbial communities consist of bacterial fungi, and protozoa etc. Therefore, understanding the diversity, pivotal role of the rumen microbes and secondary metabolites in digesting fibrous feed, providing nutrients to the host animal, defence and determining livestock host-environment interaction is key for sustainable animal husbandry. Pertinent global issues of interest include climate resilience, fight against climate change and vector borne diseases through rumen environment manipulation, to make livestock part of the solution.

The relationship between some members of the microbiome and rumen function is well known (5, 13). The role of diet on microbes diversity has been investigated (8,14,15). Whereas host genetics have been studied in determining rumen microbes (16, 17, 18), most of the studies have been done on a single species and biased towards cattle, and no comparative studies have been reported between diverse ruminant animals that vary both in feeding regime and resilience, which is the main focus of this study. Here, using four ruminant livestock that vary in feeding regime, drought resilience, and disease prevalence, we show that each livestock species created mutual association with signature microbes and secondary metabolites that provide useful ecological traits.

## RESULTS

### Distribution of bacterial and fungal populations in the rumen

To correlate the secondary metabolites with rumen microbes we performed genomic analysis of the two main rumen domains, bacteria and fungi. The taxonomic analysis of bacterial and fungal populations in the rumens of cattle, sheep, goats and camels revealed a variation in dominance of core groups of rumen microbes among the four ruminants (Fig. 1). A total of 1052 species-level, bacterial operation taxonomic units (OTUs) were uniquely identified in camels, 949 in cattle, 1065 in sheep and 847 in goats respectively (Fig. 1B). Whereas 113 bacterial (OTUs) were shared by all the four ruminants, 187 OTUs were shared by both camels and goats while 208 (OTUs) were common in cattle and sheep (Fig. 1B).

**Fig. 1:**
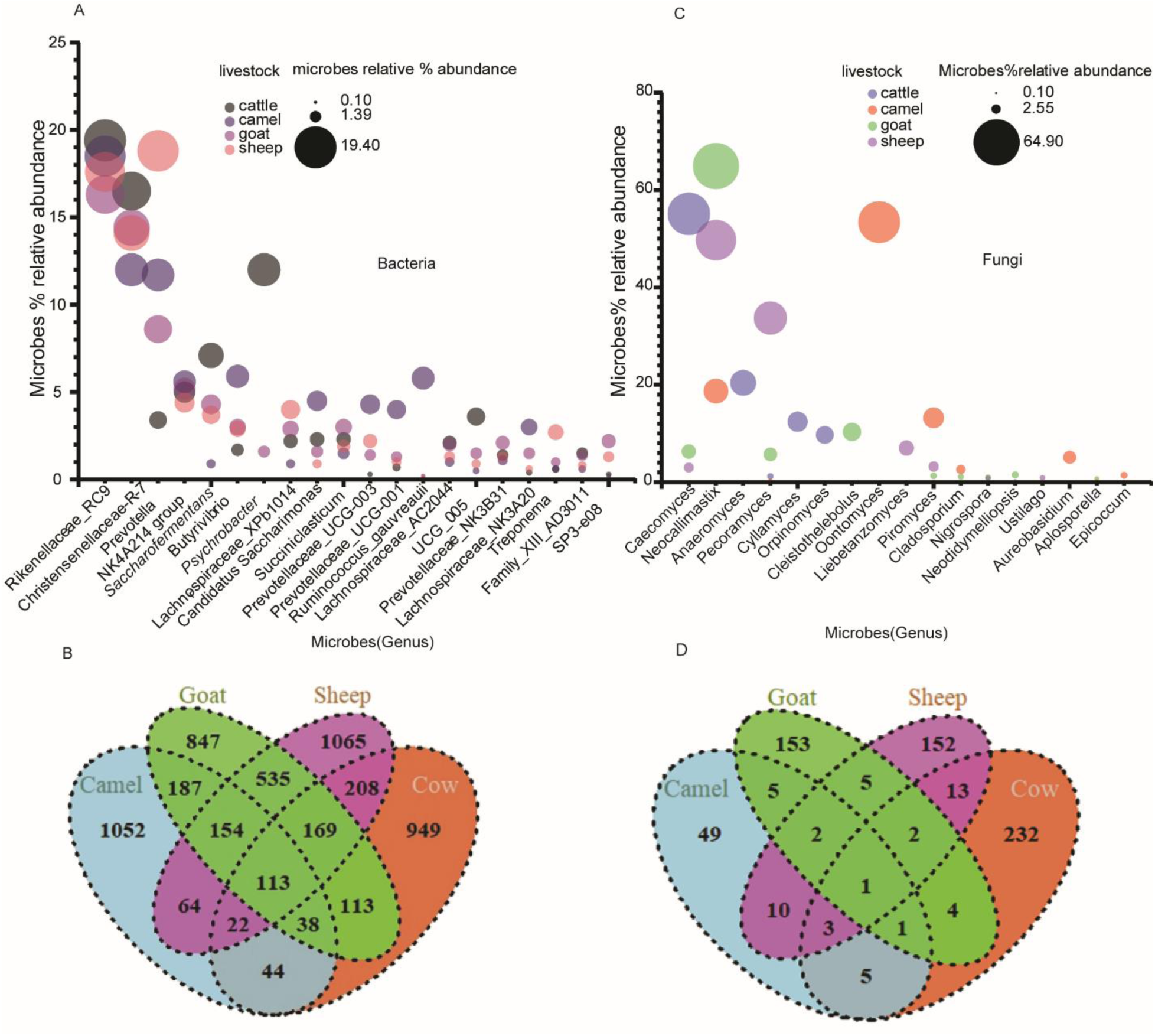
Bacteria and Fungi community compositions in different livestock A bubble plot showing the qualitative and quantitative difference of bacteria, plot generated using bacteria relative abundance data with at least 1% relative abundance in one of the four livestock species. (B). Venn diagrams showing the identified bacterial Operational Taxonomic Units (OTUs) unique and shared between livestock species (C) A bubble plot showing the qualitative and quantitative difference of fungi with at least 1% relative abundance in one of the four livestock species. (D) Fungal Operational Taxonomic Units (OTUs) among different livestock.

Bacteria being the main members of the rumen microbiome, were widely dominant across the four livestock groups analyzed, comprising most of the species richness, with some bacterial genera being livestock specific (Fig. 1A, B). A further analysis of the identified bacterial operation taxonomic units (OTUs) revealed twenty most abundant bacterial genera present among the four livestock species (Fig. 1A, C). In all four ruminants, the *Rickenellaceae RC9, Christensenellaceae R-7 group, NK4A214* group and *Succiniclasticum* group are conserved both in their presence and in their high abundance (Fig. 1A). Genus *Ruminococuce* found abundantly in camels and in small amount in goats. *Prevotella*, and *Prevotellaceae* a hydrogen-producing bacterial genus, was dominant in camels, however less abundant in cattle, goat and sheep (Fig. 1A). The *Psychrobacter* genus was found uniquely in cattle and goats but dominant in cattle but absent in sheep and camel (Fig. 1A, Table S 1). All the remaining bacterial genera were conserved in all four livestock species, but with varying abundance. Compared to camels and sheep, cattle and goats had more bacterial diversity, due to an additional genus, *Psychrobacter* (Fig. 1A, Table S 1).

A comprehensive analysis was conducted on the operational taxonomic units (OTUs) of fungi in four livestock species, namely cattle, camels, goats, and sheep. The results showed a total of 232 OTUs in cattle, 49 OTUs in camels, 153 OTUs in goats, and 152 OTUs in sheep at the genus level (Fig. 1C, D). Among these OTUs, a diverse population of seventeen highly prevalent fungal genera was found (Fig. 1C). The analysis further showed that only one fungal operation taxonomic units (OTUs) was common in all the four livestock species, while 5 were common in camels and goats whereas cattle and sheep shared 13 OTUs (Fig. 1D). Goat had the highest representation of fungal genera with camels having the least representation among the four ruminants (Fig. 1D). The anaerobic fungi genus *Caecomyces* was abundantly present in cattle, and in small amount in goats and sheep, but missing in camel. An aerobic fungus genus *Oontomyces* exclusively found only in camels with high abundance. *Neocallimastix* is the most abundant both in goats and sheep, present in camel in small amount, but missing in cattle. *Pecoramyces,* was found only in sheep and goat, in former much abundant, but missing from camel and cattle (Fig. 1C, Table S 2). *Liebetanzomyces* are only found in sheep. Furthermore, *Anaeromyces, Orpinomyces* and *Cyllamyces* were unique to cattle, whereas *Piromyces* were the major groups in camels (Fig. 1C, Table S 2). *Nigrospora* is absent in cattle, but present in the other three livestock in small amount. *Caecomyces was* dominant in cattle, absent in camel, but present in small amount in goat and sheep*. Cleistothelebolus* was distinct to goats, may be considered as signature fungal community in goat rumen (Fig. 1C, Table S2). Only *Cladosporium and Pecoramyces* are conserved among the four livestock demonstrating their requirement for conserved function. Furthermore, camel harbour different protozoans as compared to other livestock (data not shown).

### Alpha and Beta Diversity

The four ruminants showed greater variability and diversity in bacterial and fungal populations as revealed by Shannon, Chao1 and Pielou evenness, alpha diversity indices (Fig. 3A-F). However, cattle and goats showed similarity in evenness and richness. Beta diversity was assessed by calculating the PCoA of different rumen bacterial and fungal domains using Bray-Curtis method. The analysis revealed significant dissimilarities in bacterial and fungal domains distributions among ruminants as displayed by the different eclipse clusters (p = 0.004, PERMANOVA; Fig. 2G-H). Compared to other ruminants, cattle and camels showed higher variability of bacterial communities (Fig. 2G), whereas camels exhibited a different fungal domain population (Fig. 2H).

**Fig. 2:**
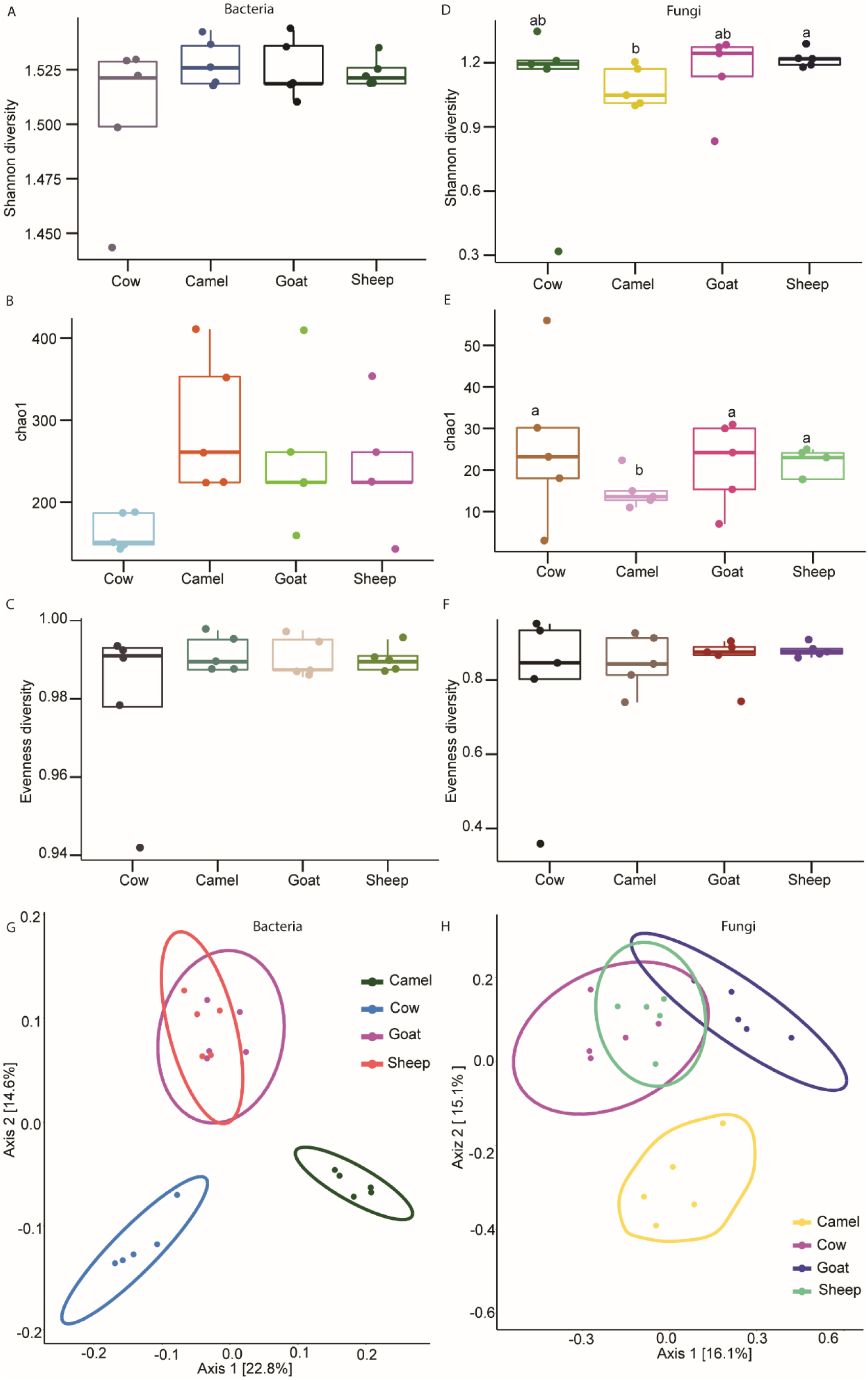
Alpha (A-F) diversity indices for bacterial and fungal populations using Shannon index (p = 0.77, and 0.28; A and D). Chao1 richness index estimates of bacteria and fungi (p =, 0.036, 0.24; B and E). Evenness estimates in bacteria and fungi (p = 0.87, 0.95; C and F). Beta diversity PCoA ellipse clusters based on unweighted unifrac distance dissimilarity method showing the distribution of bacteria and fungi (G-H). Bars followed by different letters are statistically significant.

**Fig. 3:**
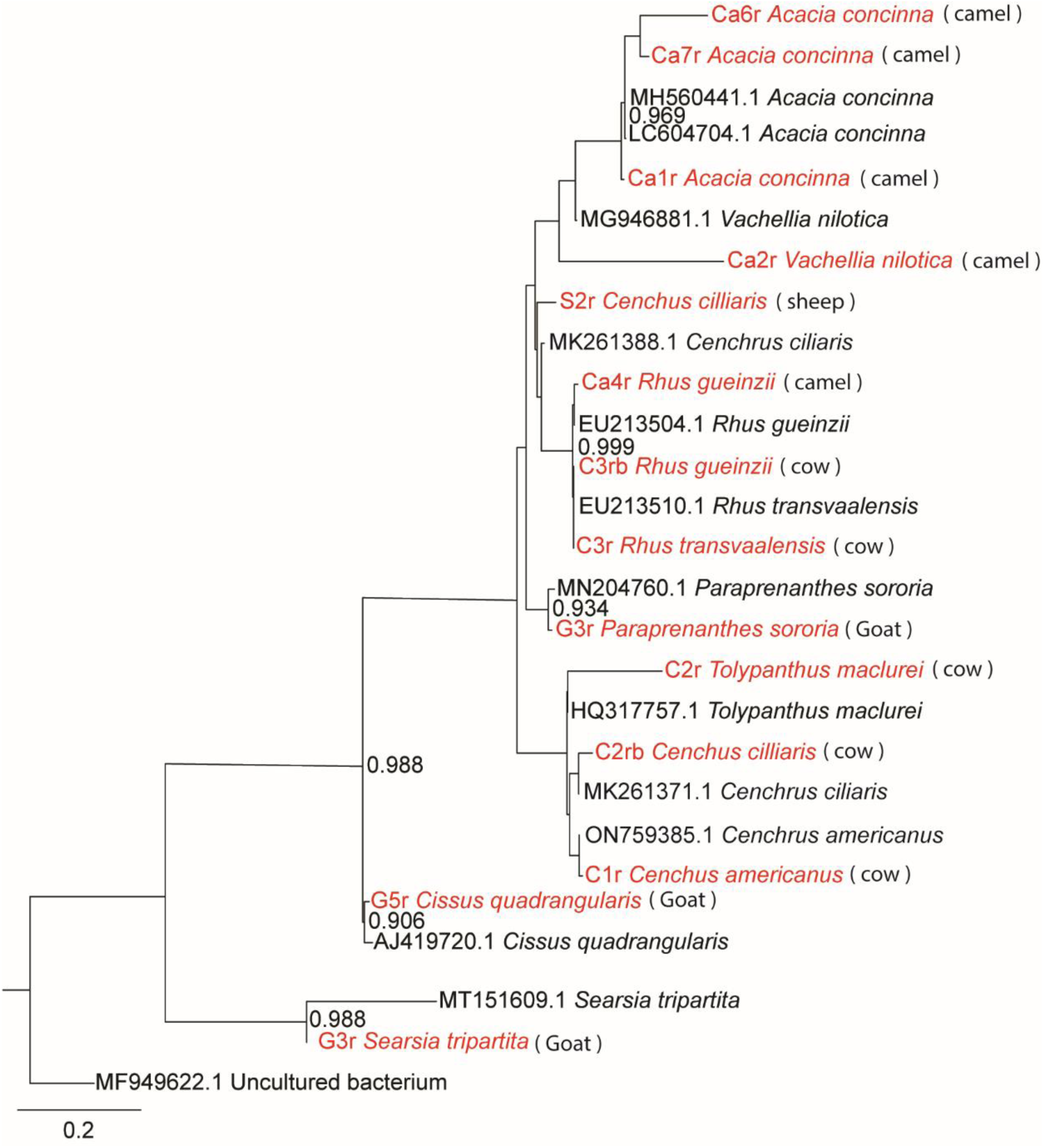
Phylogenetic tree showing plant diet composition identified in livestock rumens, analysis based on maximum likelihood, branch values indicate bootstrap support of 1000 pseudo replicates.

### Dietary composition assessment in livestock rumen

Microbial diversity and secondary metabolites may be affected by host diet composition beside host’s individual genetic makeup (8). To understand the overall metabolites make up in relation to diet among ruminants, the study characterized the various diet consumed by the four livestock. In pastoralist setup where feed is not controlled or restricted, livestock can feed on a wide range of plant materials. For instance, we found that in addition to grasses (*poaceace*), *Cenchus cilliaris* and *Cenchus americanus,* which had been consumed by cattle and sheep, cattle had consumed other plant species such as *Rhus gueinzii and Rhus transvaalensii* despite being predominantly grazers (Fig. 3). Unlike cattle and sheep, camels and goat are specially adapted to feed on leaves, fruits of high-growing woody plants, soft shoots and shrubs, such as *Acacia concinna*, *Paraprenanthes sororia*, *Vachellia nilotica* and *Searsia tripartita* (Fig. 3), which are predominantly found within arid and semi-arid areas. Therefore, points to the diversity in dietary composition among the ruminants, which influences both the metabolite compound and microbial population composition among the ruminants.

### Ruminal metabolite composition in livestock

In the present study, a total of 162 metabolite compounds (Table S 3) were identified in bovine rumen content of four livestock; cattle, sheep, goat and camel by GC-MS. The detected compounds represented various chemical classes, including alcohols, ketone, phenols, volatile fatty acids, terpenes, esters and hydrocarbons. Although most major classes of secondary metabolites have ubiquitous distributions among the four livestock, each livestock species has its own signature secondary metabolites (Fig. 4A). A random forest classification was conducted to reveal the top 10 predictive compounds for individual ruminant species (Fig. 4B). 2, 6 dimethyl, 4-octene, 3 Methyl, butanoic acid, 1, 3 cyclohexene and tricyclene being the most predictive secondary metabolite compounds of cattle, camels, goats and sheep rumen, respectively. In camel 5 out of the ten predictive compounds are acids, signifying the diversity of acid in camel rumen. However, we did not observe dominance of any specific chemical class in the other livestock, diverse class of compounds contribute for predictive signature odours. The diversity and contrast in metabolite composition among the ruminants was revealed by the clustering and segregation of the respective species based on their metabolite composition by multidimensional scaling (MDS) and matrix plot (Fig. 5A-B). While some species such as cattle, and sheep, which are grazers clustered in close proximity, camels and goats, which are browsers, were distinctly clustered apart from the other ruminants and also from each other (Fig. 5B). Thus, demonstrating a similarity in metabolite composition among grazers (cattle and sheep) and but not clear with browsers (camels and goats). Overall, the four livestock dissimilarity based on their rumen secondary metabolites was 72.5%. Twenty-three compounds, based on quantitative and qualitative difference contributed for more than 50% of the variation (Fig. 5C). Acids are found in most abundant in camel, the three livestock produced almost 2x carbon dioxide as compared to camel. Isoamyl benzyl ether absent in cattle, linalool propionate detected only in sheep, valence detected only in cattle, citronellene-beta absent in cattle, cis-calamine absent in camel, menthane-1-p absent in cattle and sheep, 1, 3, cyclohexadiene found only in goat and sheep, terpinolene found only in sheep, β-gurjunene absent in camel and goat, limonene absent in cattle, 1, 5, 9-undecatriene absent in camel.

**Fig. 4:**
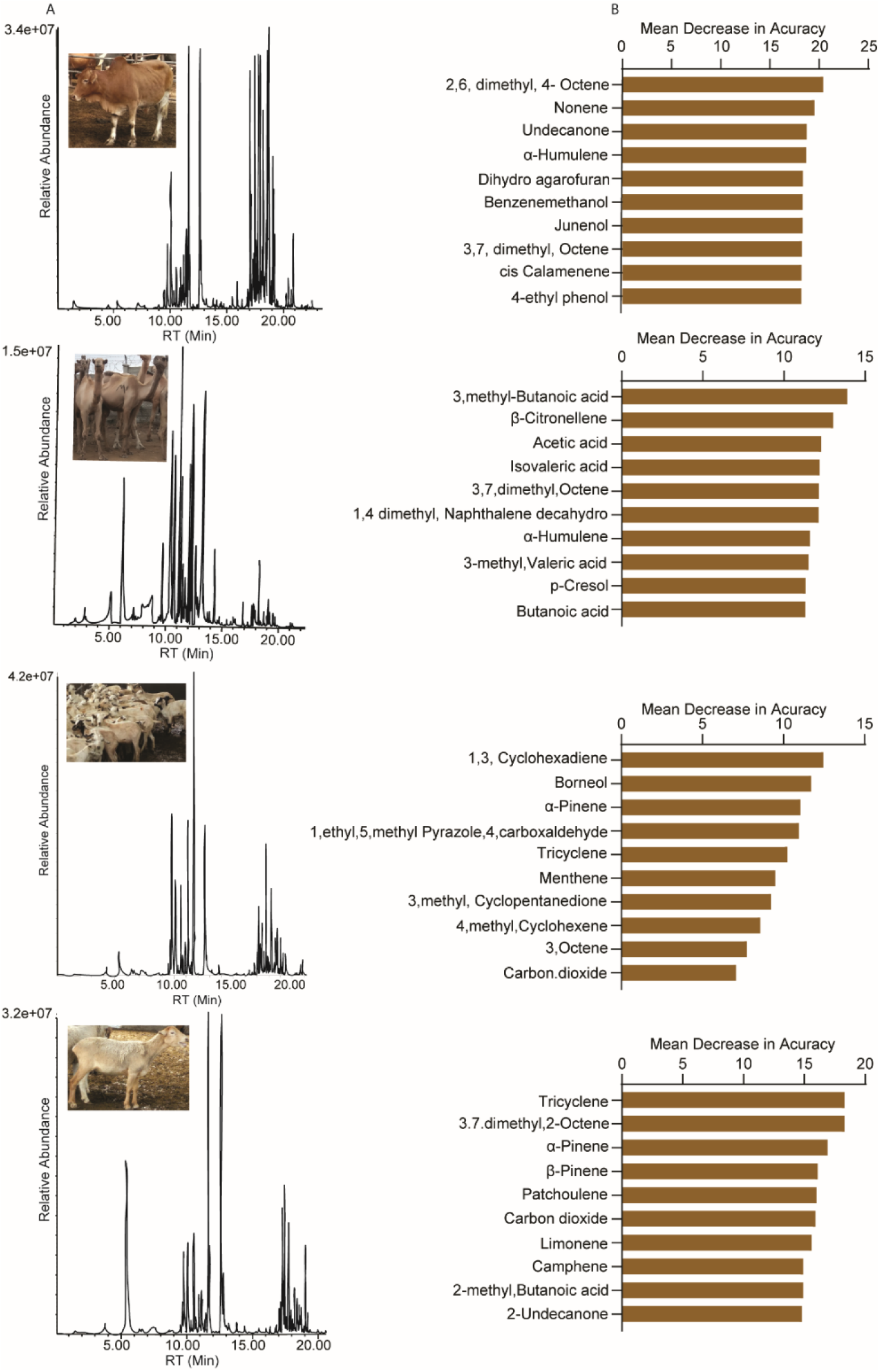
GC-MS chromatogram profiles of metabolite compounds in ruminal fluid of various livestock (A), cattle, camel, sheep and goat respectively. Histograms showing the classification of the top ten predictive compounds from different livestock rumens based on their Mean Decrease in accuracy (MDA) of the Random Forest analysis (B).

**Fig. 5:**
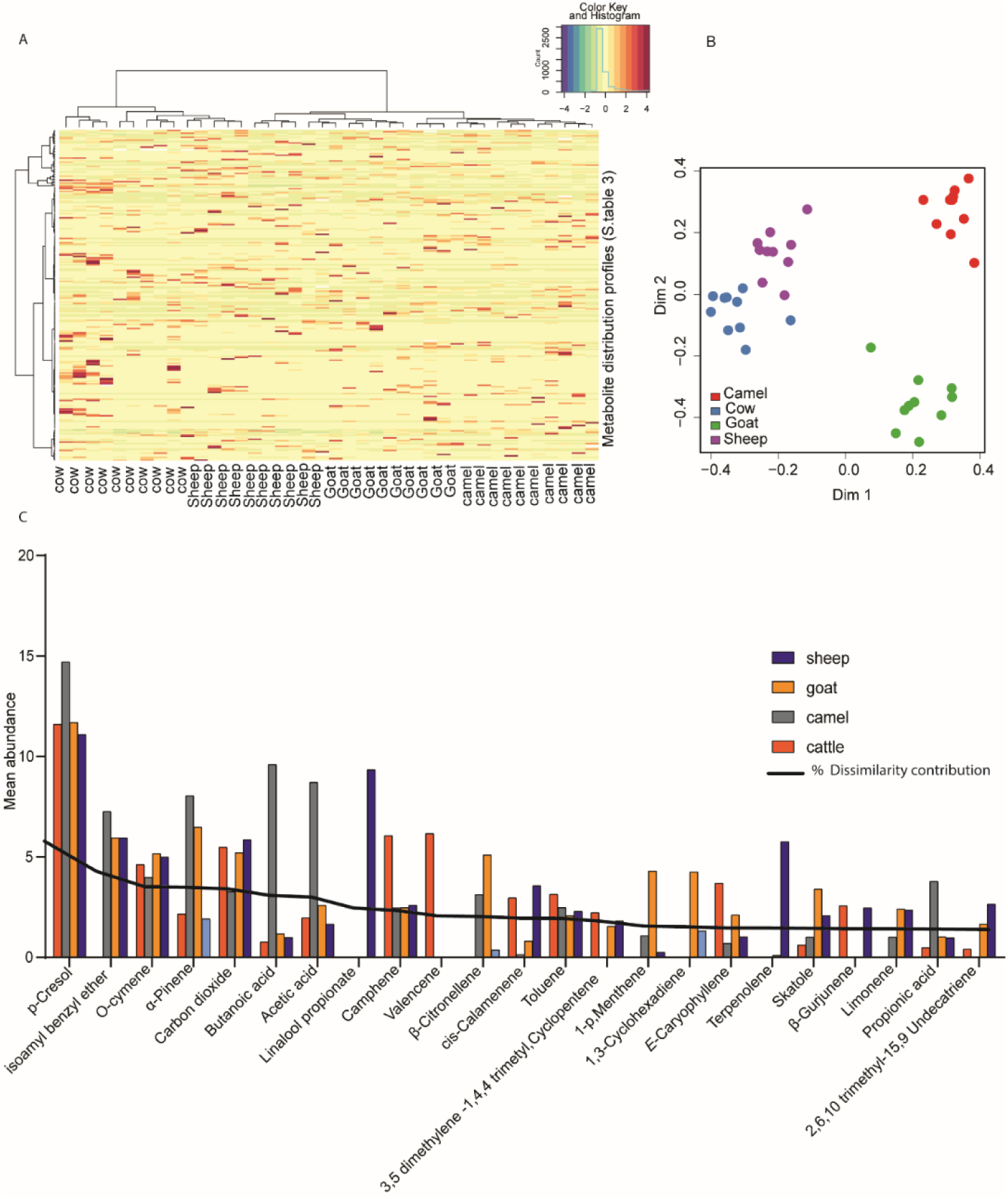
Heatmap coded matrix showing relative percent contribution of individual compounds to the total composition of each livestock species (A) (for detail please see (Table S 3). Multidimensional scaling (MDS) plot showing the segregation of ruminants based on metabolite composition (B). Histogram showing the classification of the top twenty-three for 50% dissimilarity contributing metabolite compounds from all livestock based on SIMPER analysis, the line graph shows the percentage contribution of a given compound for the dissimilarity. (C).

### Variability in metabolite composition among individual species population

The correlation between populations of the same species dynamics and the metabolite compound profiles of four types of livestock was investigated using a Pearson’s correlation analysis. Generally, a minimal variability in metabolites was observed between individual of the same species. Cattle, goats, and camels showed minimal variability in their volatile organic compound profiles (Fig. 6C); however, sheep populations showed some variation in sheep 7, 8, 9, and 10 (Fig. 6D). As a result, the rumen odor profiles in the herd populations of the four cattle species used in this investigation were comparable.

**Fig. 6:**
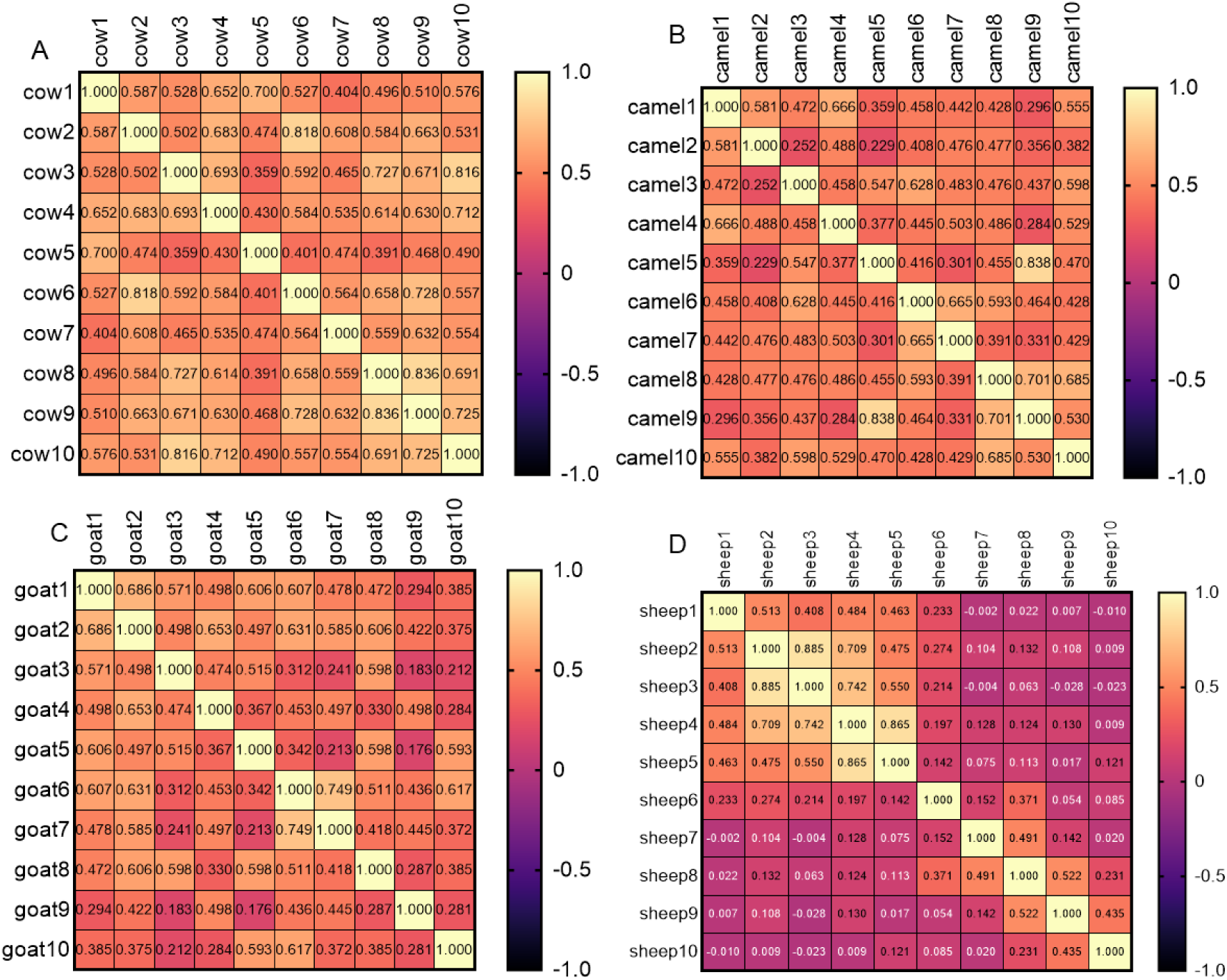
Color coded Pearson’s correlation plots for identified volatile organic compounds among individual animal in their respective species.

### Metabolite composition by chemical functional groups

We then evaluated the chemical identities and variation in distribution of volatile organic compounds across the four ruminant species, with compounds categorized based on their functional group classification, including phenols, alcohols, indoles, monoterpenes, sesquiterpenes, acids, hydrocarbons, and ketones. We found significant differences in the relative abundance of certain chemical class, such as alcohols, hydrocarbons, monoterpenes, acids, and sesquiterpenes, among the four livestock groups, with cattle, sheep, camel, and goats displaying varying relative abundance of these compounds (Fig. 7A-G, P < 0.05). Cattle, sheep, and camel had significantly lower alcohols and sesquiterpenes concentrations compared to goats (Fig. 7A & E, ANOVA, P < 0.05). Similarly, acids varied between the four ruminants, with camels having significantly high acid abundance compared to cattle, sheep and goats (ANOVA, P = 0.004) (Fig. 7B), that resulted in acidic rumen environment in camel. A significant variation was noted in camel ruminal pH compared to the other three livestock groups (ANOVA, P < 0.05). Cattle, goat and sheep had a relatively neutral pH ranging from (7.0 -7.4), compared to camel which had a acidic pH (pH 6.3-6.5). Phenols, ketones, and hydrocarbons are more abundant in goats compared to the other livestock, sheep has more indoles compared to other livestock (ANOVA, P< 0.05) (Fig. 7H).

**Fig. 7:**
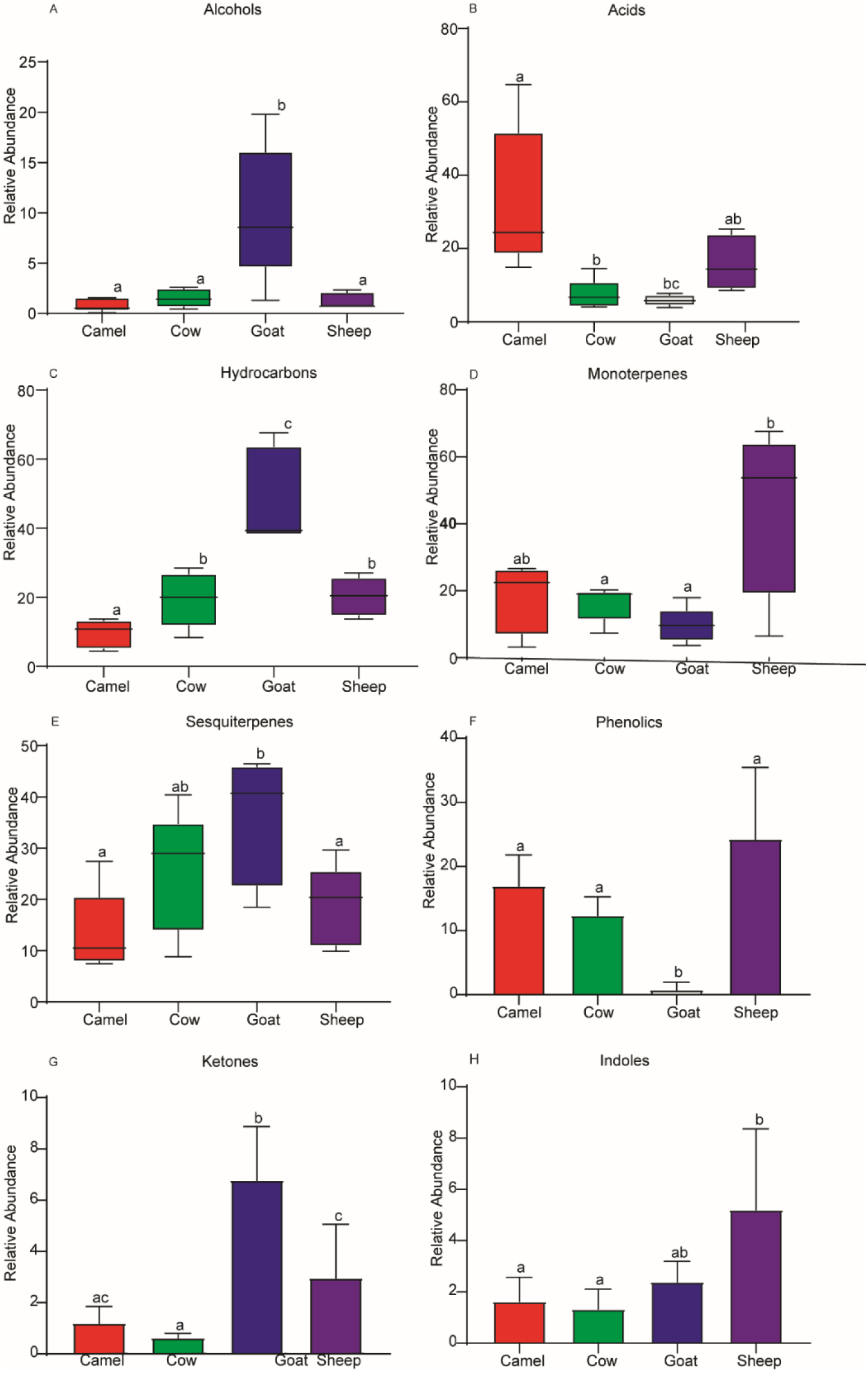
Box plots showing variation of chemical families of identified metabolite compounds among different ruminants (A-H). Bar graphs followed by different letters are statistically different in their abundance.

## DISCUSSION

In this comparative study we show that there is a complex network of rumen microorganisms that coexist, interact and compete for substrates resulting in a critical balance of various end products such as secondary metabolites without autotoxicity to provide energy for microbial growth, and beneficial end products for the host. A better understanding of how the rumen microbiome influences host health and performance may lead to novel strategies and treatments for trait improvement in the livestock sector in nature inspired ways.

Three bacterial genera (*Rikenellaceae RC9 gut group, Prevotella, NK4A214 group and Christensenellaceae R-7 group)* were highly conserved both in their presence and abundance among the four livestock species regardless of genetics which may suggest that they are core rumen bacteria, essential for highly conserved common traits or function for these animals. However, there were unique bacteria in a given host, for instance *Anaeromyces, Cyllamyces and Orpinomyces* found only in cattle, which may be suggestive of the existence of species specific microbes that requires host rumen-microbes’ environment balance. Similarly, two fungi genera *Cladosporium and Pecoramyces* are conserved between the four livestock in their presence, but not in abundance demonstrating they are necessary for a conserved function, and their function may be relative abundance dependent in each species. Several fungi genera are unique either present only in one animal species or shared between some of them but not by all the four, even those that are shared present in different abundance, that may be integral to their environment and potentially compatible with rumen environment and host requirements. The significant variation in fungi microbes between the four rumen livestock may be explained to a significant extent by host genetics (5, 16, 17, 18). This unique microbes-host framework variation in microbial composition between hosts may affect microbially-mediated ecosystem processes as well as may depend on host phylogenetic relatedness and trait-based patterns of ecologies (19). Each of the bacterial and fungal communities established in the present study, play a specific metabolic role in the rumen (20, 21, 22). For instance, bacterial species like *Ruminococcus, Lachnospiraceae, Christensenellaceae* and *Prevotella* are associated with hydrogen production during rumen fermentation (23, 24). There are some microbes that were ubiquitous in all four rumen species demonstrating their wide rumen environment adaptation, for instance camel rumen has acidic pH compared to other livestock.

The next step is to explicitly link the observed microbial, secondary metabolites diversity and network with basic evolutionary principles, which is biological fitness. We have shown the variation between diverse microbes among the four livestock that varies in feeding behaviour, draught resilience and disease susceptibility (25). For instance, camel and small ruminant to some extent are the most resilient among analyzed livestock to frequent drought as compared to cattle (9,10,11,12). This could be due to the abundantly presence of unique anaerobic fungi, *Oontomyces*, originally identified from Indian camel (26) and bacteria (*Prevotella)* in camel that have demonstrated high capability of diet conversion (27, 28, 29, 15). Additionally, fungus *Neocallimastix,* which is present in camel, sheep and goat but absent in cattle, have been shown to be effective in bioconversion potential of poor diet such as lignocellulose into useful products (30, 31) may have contributed for their resilience. These microbes combine with other physiological mechanism such as suppression of cholesterol biosynthesis in kidney of camel to retain and reabsorption of water (32) may contributed for camel draught resilience. Thus, camel’s evolutionary success to dry climate, partly may be due to the ability to engage in mutualistic interactions with useful microbes that provide novel ecological adaptation traits. Furthermore, such knowledge will give us opportunity to manipulate rumen environment to make livestock less susceptible to vectors, efficient in converting their diet to animal protein and to make livestock environmentally friendly. For instance, in one study, the addition of a fungal inoculant to the diet of dairy cows was found to increase the production of propionate and decrease the production of acetate (33), which is a precursors of greenhouse gas production. Furthermore, microbiome work in humans and rodents has revealed that microbes play essential roles in host health and function (34, 35). Similarly, in our previous work these various livestock exhibited various susceptibility to various pathogens (36) that may depend on their mutualistic association with useful microbes.

The study conducted by (5) shown that the core microbiome had a significant explanatory role in relation to dietary components within a controlled experimental setting. In our experiment, it was difficult to dissect the role of diet for the microbes and chemodiversity variation as the animals were from free grazing and browsing set up and fed on diverse diets. If we assume that diet may structure rumen microbes we would have expected similarity both in microbes and secondary metabolites between browsers (camel and goat) and also between grazers (cattle and sheep). However, we did not find a clear link between diet and microbes, for instance only one bacterial genera *Psychrobacter* is missing in camel and sheep. If diet shape the rumen microbes, browsers (camel and goat) should share more similar microbes than camel has in common with cow and sheep and vice versa. On the other hand, if microbes dictate diet, cattle and sheep share more similar microbes than what camel and cattle share. But sheep and goat shared more bacteria than either of them shared between camel and cattle. This may also be because there is no strict browsers and grazers under free grazing setting as they can easily shift between various diets depending on feed availability. The various plants consumed by the various livestock are characterized by high fiber content, and rich in secondary metabolites, and bioactive compounds including tannins, flavonoids, alkaloids, and terpenoids which may have potential health benefits for ruminants (37, 38, 39). The utilization of shrubs and woody plants in livestock diets has been shown to increase rumen metabolite richness compared to diets based on traditional forage sources (40, 41, 42). Studies showed that feeding goats on *Acacia saligna*, a shrub species, led to increased diversity and richness of rumen metabolites compared to a control diet based on alfalfa hay (40, 41, 42, 43). The composition of the plant diet can have significant impacts on the production of metabolites in the rumen for instance, Grasses (*Poaceae*) contain fermentable cellulose, hemicellulose, lignin, and protein which are broken down by rumen microbes into various metabolites, including acetate, propionate (13, 44), which are ingredients in greenhouse gas formation and energy source. Hence the variability of plant diets can have a significant impact on rumen metabolite production and composition in livestock.

Rumen fermentation is a complex process that results in the production of various metabolites (5, 33). We established a wide range of secondary metabolite compounds in rumen, which is an interplay between, host genetics, diet and microbes, most of which are associated with various biochemical activities in livestock rumen. The detection of metabolite compound classes, such as volatile fatty acids, aromatic hydrocarbons, terpenes, hydrocarbons, phenols, and alcohols, displays the diversity and complexity of metabolic synthetic pathways in livestock rumens, leading to the production of several diverse metabolites (45). The detection of plant-derived metabolite compounds such as camphene, α-pinene, and β-caryophyllene, including fecal predictive indolic and phenolic compounds like p-cresol (a byproduct of protein breakdown in animal gut) and skatole, which had previously been reported in various animals metabolic by products, for instances in animal feces, which have role in livestock-vectors interaction (45, 46, 47, 48), demonstrate that metabolites are conserved as they pass through various digestion process. But we also observed less complexity in some metabolites for instance, phenols in the rumen are less complex as compared to livestock urine (36), demonstrating metabolites may gain complexity after they left rumen.

Even though the examined metabolite composition varied among the ruminants, minimal intraspecific variation was realized among individual species herd, indicating a potential host specific microbes and host genotype effect (49) implies that rumen secondary metabolites may not be affected by livestock population dynamics as was demonstrated by (36, 50). Despite their diversity among the 4 ruminants, identical metabolite compound classes were detected from rumen metabolism, point to a similarity in their biosynthetic pathways between the four livestock species and those metabolites may have conserved function regardless of the host genetics.

Studies have highlighted a direct relationship between bacterial and fungal populace with rumen metabolome (51, 52, 53, 54, 55, 56). These microorganisms work together in a symbiotic relationship with the host to break down complex plant polysaccharides and fiber into simple sugars, which can then be fermented into volatile fatty acids (VFAs), microbial proteins and other metabolites that can be absorbed by the host animal (8, 57). Thus, the various secondary metabolites identified may provide various functions to the host. Ruminant, such as cattle, sheep, and goats utilize hydrocarbons as energy source largely contained in plant carbohydrates like glucose and sucrose, by fermenting them in their rumen into volatile fatty acids, which are then absorbed and utilized for energy (58, 59). In this study, we established notable differences in hydrocarbons, terpenes, ketones, and indoles relative abundance among the ruminants. Such variations clarify relevant aspects such as diet composition, breed, and environment, since the detection and concentration of most ruminal metabolite compounds are influenced by these factors (60, 61, 62). The diversity and importance of different compound classes of rumen metabolome in livestock were further demonstrated by the detection of terpenes, which have been linked to improve nutrient utilization and digestive health (63). In addition to terpenes, chemical compounds classes like acids, phenols, indoles, ketones, and alcohols, also varied significantly among the ruminants (Fig. 7).

Acids profile significantly differs between the four livestock species, being the highest in camel. Acids are involved in the hydrolysis of complex carbohydrates, such as cellulose, lignin and hemicellulose, into simpler sugars that can be further metabolized by rumen microbes (64). This may be ascribed to the fact that acids are energy sources for the host animal and can be used as precursors for energy production during special conditions. For instance, fatty acids such as acetate is used by the host animal as a precursor for fatty acid synthesis in adipose tissues, which can then be utilized as an energy source during times of high energy demand, such as during lactation or periods of feed restriction (65), thus the diverse acids produced in camel rumen may have contributed to camel rumen acidic pH, resilience even during extended drought, with limited feed availability in arid and semi-arid ecologies. Phenols and indoles are aromatic compounds that are derived from lignin, which is present in the cell wall of plants, are produced during the fermentation of plant material in the rumen, and have been shown to have antimicrobial properties that can help to maintain a healthy microbial balance in the rumen (66), and antioxidants, which can help to reduce oxidative stress in the rumen and improve animal health (67, 68). Alcohols, provide energy for rumen microbes in addition to being a carbon source for the synthesis of microbial protein (69) and ketones shown to be an alternative energy source for ruminants in addition to preventing ketosis (70). Furthermore, elucidation of maternal, genetic, and environmental factors, rumen environment (for instance pH, nutrient etc) that influence rumen microbiome establishment and development may provide novel insights into possible mechanisms for manipulating the rumen microbial and secondary metabolites composition to enhance long-term host health, performance and climate resilience.

## Conclusion

We have documented various microbes and secondary metabolites which vary among rumens that may provide, useful traits, such as energy source, antioxidant, digestive and detoxifying capabilities, improve host defense against pathogens. We can conclude the diversity both qualitative and quantitative in microbes may contributes to the variation observed between the four livestock phenotypic traits expressed by the host animal including chemodiversity and resilience. Our result may have application in rumen environment manipulation targeting microbes and secondary metabolites network to make livestock productive, resilient, and less susceptible to vectors and environmentally preferred, climate smart livestock husbandry. Our results demonstrate rumen fermentation at the interface of host genetics, microbes and diets has a significant implication for the production of complex secondary metabolites, which in turn can confer unique ecological traits to the host organisms.

## MATERIALS AND METHODS

### Collection of rumen content

Bovine rumen contents were collected from 10 different freshly slaughtered boran cattle (*Bos indicus*), goats (*Capra aegagrus hircus*), sheep (*Ovis aries*) and camels (*Camelus dromedaries*) from their respective abattoirs in Nairobi and Machakos County, respectively. The samples (500ml each) were kept in sterile airtight freeze-resistant 1L odor collection glass jars (Sigma Scientific, USA) and transported in a cooler box to the laboratory for metabolite compound collection and analysis.

### Genomic DNA extraction

To extract genomic DNA from rumen contents of cattle, sheep, camels, and goats, 200 μl of the sample was mixed with an equal volume of buffered phenol and 20 μl of 20% SDS in a 2 mL centrifuge tube (Eppendorf, Germany). After adding 0.5g of 2 mm zirconia beads (BioSpec Inc., USA), the mixture was shaken thrice in a mini-tissue lyser (Qiagen, Hilden, Germany), at a frequency of 30Hz for 90 seconds. The lysate was then centrifuged at 14000 rpm on a 5417R centrifuge (Eppendorf, Germany) for 10 minutes, and the supernatant was transferred to a 1.5ml clean tube (Eppendorf, Germany). Afterwards, 200 μl of buffered phenol was added to the supernatant, the mixture was briefly vortexed, and then centrifuged at 14000 rpm at 4°C for 15 minutes. The DNA was then precipitated by adding 500 μl absolute ethanol to the supernatant in a clean 1.5ml centrifuge tube and centrifuged at 14000 rpm at 4°C for 5 minutes. The supernatant was discarded, and DNA pellet washed by 500ul of 70% ethanol then centrifuged for 5 minutes. Finally, the pellet was suspended in 100 μl of preheated elution buffer G (ISOLATE II Genomic DNA kit, Bioline Meridian). The DNA quality and quantity was checked by Nanodrop spectrophotometer (Thermo Scientific, Wilmington, DE, United States). Aliquots of 50μl of the obtained DNA extracts were sent to Macrogen Inc (Netherlands) for Illumina next-generation sequencing (NGS) targeting 16S rRNA and ITS1 for bacteria and fungi respectively. The remaining amounts (50μl) were utilized for PCR for plant diet identification.

### PCR amplification for diet composition screening

PCR amplification targeting two chloroplast markers, consisting of coding (rbcL gene) and non-coding gene spacer region (trnH-psbA) primers (Table S 4) was done according (71). The obtained amplicons were then sent for sequencing at Macrogen Inc (Netherlands). Using Geneious software, obtained sequences were cleaned, edited, and aligned, resulting in a congruent sequence made up of contigs from both the forward and reverse sequences. The plant species were then identified by aligning the processed sequences against the GenBank database using the NCBI BLAST1 search engine. Subsequent phylogenetic analyses was done using the MEGA software version 11 (72).

### Metabolite extraction and analysis

Metabolite compounds from cattle, camel, sheep, and goat rumen contents were extracted using the headspace, solid Phase microextraction (HS-SPME) technique as detailed by (73). Stableflex 24Ga, manual holder SPME fibers (65μm, Polydimethyl Siloxane/Divinylbezene (PDMS/DVB), Supelco, Bellefonte, Pennsylvania, USA) were used to trap the volatile metabolite compounds, and later analyzed by Gas chromatography (GC, HP-7890A, Agilent technologies, USA) coupled with Mass Spectrometry (MS, 5975C, Agilent technologies, USA), after which the compounds were identified as described by (73).

### Data analysis

Multivariate statistical analyses were conducted based on the nature of the obtained data using R studio statistical software version 4.2.1 (74), PAST software Version 4.03 (75) and GraphPad Prism version 9. Similarity percentages (SIMPER) and One-way ANOSIM with Bray-Curtis dissimilarity index was used compare the profiles, and establish dissimilarity contribution of identified metabolite compounds based on their peak areas across the four livestock species. The metabolite compounds were then classified using the R software package “Random Forest”, version 4.2.1. The random forest analysis was executed by running 1000 iterations (ntree) with 10 compounds randomly selected at each split (mtry=√q, where q is the total number of compounds. Based on the function ‘importance ()” we generated the mean decrease in accuracy (MDA), which provides an importance score for each metabolite compound. For each livestock, the metabolite with the highest MDA value was considered the most important. A multidimensional scaling plot (MDS) and a classical cluster dendrogram were used to visualize the output of analyzed metabolite compound profiles in each livestock. We then used Pearson’s correlation to establish how metabolite compounds compared among individual ruminants’ herd population. The detected metabolite compounds from across the 4 ruminants, were then pooled based on their chemical identities, after checking for normality using Shapiro-Wilk test (P > 0.05), Pairwise comparison of the mean relative abundance of respective metabolite compounds in each chemical entity was analyzed by analysis of variance (ANOVA) among the four ruminants. Statistical significance was declared at P < 0.05.

### Bioinformatics analysis

Initially, the data obtained from Illumina sequencing was assessed using <u>nf-core-ampliseq</u> (v2.4.0) workflow and nextflow (v22.10.0), with predefined parameters of trunclenf = 180 and trunclenr = 120. The workflow proceeded as follows: first the quality of the reads were checked, using FASTQC (version 0.11.9). Cut adapt (v4.1) was then employed to trim reads and eliminate adapter sequences, following the method developed by Marcel Martin (76) . Preprocessing was performed using the DADA2 tool (v1.26.0) for filtering and trimming, dereplication, sample inference, merging of paired end reads, removal of chimeras, and taxonomic classification of the ASVs, as outlined by Callahan et al. (2016). Furthermore, DADA2 performed the classification of the ASVs’ taxa based on their taxonomic categorization (Silva database v138 was used on 16S rRNA, and unite database v8.3 was used on ITS1 rRNA). Finally, Barrnap tool (v0.9) was employed to predict the location of ribosomal RNA genes in genomes.

### Abundance Visualization

To visualize the ASV count table and ASV taxonomy table generated by the DADA2 algorithm within the nf-core ampliseq workflow, R statistical software (version 4.2.1) was used for further analysis. The ASV count table, ASV taxonomy table, and metadata were into a single phyloseq object using the Phyloseq package (version 1.40.0) in R. A *subset_taxa()* function was then employed to eliminate undesired taxa before converting it to a data frame for further manipulation using the *phyloseq_to_df()* function. Subsequent data frame manipulation was conducted by tidyverse package (version 1.3.2). Lastly, ggplot2 (version 3.4.0) and Cairo (version 1.6.0) were used to produce the visual plots.

## ACKNOWLEDGEMENT

The authors gratefully acknowledge the financial support for this research by the following organizations and agencies: Max Planck Institute *icipe* partner group to MNG and the Swedish International Development Cooperation Agency (Sida); the Swiss Agency for Development and Cooperation (SDC); the Federal Democratic Republic of Ethiopia; and the Government of the Republic of Kenya. The views expressed herein do not necessarily reflect the official opinion of the donors.

We thank Mr. John Ngiela and Mr. Joseck Otiwi for their valuable support during sample collection. Mr. Onesmus Wanyama, Mr. JohnMark Makwatta and Mr. James Kabii for their guidance on chemical and molecular biology instrumentation, respectively.

## AUTHOR CONTRIBUTION

VOO, designed, collected data, analyzed data, wrote the manuscript, MNG conceptualized, designed, analyzed data, wrote the manuscript, resource mobilization. CK, SM, and NVO contributed in the bioinformatics data analysis part of the work. GBO and JMO supervised, reviewed and edited the manuscript.

The authors declare no competing interests.

## FUNDING

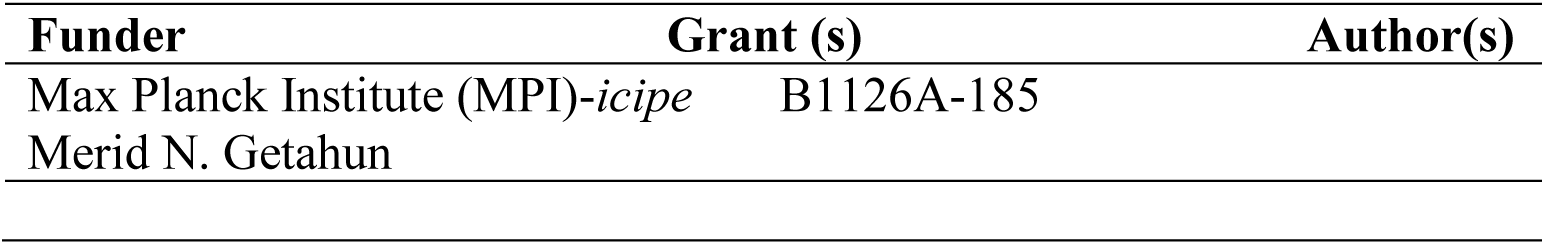

## DATA AVAILABILITY STATEMENT

The datasets generated during and/or analyzed during this study are all included in the manuscript and as supplementary materials.

### Supplemental Material

**Table S 1 Bacteria population abundance**

**Table S 2 Fungi population abundance**

**Table S 3 List of identified metabolite compounds**

**Table S 4 Primer list for diet screening**

## REFERENCE

1. Fao, I., & UNICEF. (2021). WFP and WHO. 2020. The State of Food Security and Nutrition in the World 2020. Transforming food systems for affordable healthy diets. Rome, FAO.

2. Graham, M. W., Butterbach-Bahl, K., du Doit, C. J. L., Korir, D., Leitner, S., Merbold, L., Mwape, A., Ndung’u, P. W., Pelster, D. E., Rufino, M. C., van der Weerden, T., Wilkes, A., & Arndt, C. (2022). Research Progress on Greenhouse Gas Emissions From Livestock in Sub-Saharan Africa Falls Short of National Inventory Ambitions. Frontiers in Soil Science, 2, 927452.

3. García-Yuste, S. (2020). Sustainable and Environmentally Friendly Dairy Farms. Springer International Publishing.

4. Tapio, I., Snelling, T. J., Strozzi, F., & Wallace, R. J. (2017). The ruminal microbiome associated with methane emissions from ruminant livestock. Journal of Animal Science and Biotechnology, 8(1), 7.

5. Wallace, R. J., Sasson, G., Garnsworthy, P. C., Tapio, I., Gregson, E., Bani, P., Huhtanen, P., Bayat, A. R., Strozzi, F., Biscarini, F., Snelling, T. J., Saunders, N., Potterton, S. L., Craigon, J., Minuti, A., Trevisi, E., Callegari, M. L., Cappelli, F. P., Cabezas-Garcia, E. H., … Mizrahi, I. (2019). A heritable subset of the core rumen microbiome dictates dairy cow productivity and emissions. Science Advances, 5(7), eaav8391.

6. Medjekal, S., & Ghadbane, M. (2021). Sheep digestive physiology and constituents of feeds. In Sheep Farming-An Approach to Feed, Growth and Health. IntechOpen.

7. Zeng, Y. (2017). Microbial community compositions in the gastrointestinal tract of Chinese Mongolian sheep using Illumina miseq sequencing revealed high microbial diversity. 10.

8. Zhang, Y. K., Zhang, X. X., Li, F. D., Li, C., Li, G. Z., Zhang, D. Y., Song, Q. Z., Li, X. L., Zhao, Y., & Wang, W. M. (2021). Characterization of the rumen microbiota and its relationship with residual feed intake in sheep. Animal, 100161.

9. Toulmin, C. (2009). Securing land and property rights in sub-Saharan Africa: The role of local institutions. Land use policy, 26(1), 10–19.

10. Boru, D., M. Schwartz, M. Kam and A. A. Degen, 2014: Cattle reduction and livestock diversification among Borana pastoralists in Southern Ethiopia. Nomadic Peoples, 18, 115–145.

11. Kagunyu, A. W. and J. Wanjohi, 2014: Camel rearing replacing cattle production among the Borana community in Isiolo County of Northern Kenya, as climate variability bites. Pastoralism, 4, 13.

12. Watson, E. E., Kochore, H. H., & Dabasso, B. H. (2016). Camels and climate resilience: adaptation in northern Kenya. Human Ecology, 44(6), 701–713.

13. Morgavi, E. Rathahao-Paris, M. Popova, J. Boccard, K. F. Nielsen, H. Boudra, Rumen microbial communities influence metabolic phenotypes in lambs. Front. Microbiol. 6, 1060 (2015).

14. Jang, S. Y., Kim, E. K., Park, J. H., Oh, M. R., Tang, Y. J., Ding, Y. L., Seong, H. J., Kim, W. H., Yun, Y. S., & Moon, S. H. (2017). Effects of physically effective neutral detergent fiber content on dry matter intake, digestibility, and chewing activity in Korean native goats (Capra hircus coreanae) fed with total mixed ration. Asian-Australasian Journal of Animal Sciences, 30(10), 1405–1409.

15. Jami, E., Israel, A., Kotser, A., & Mizrahi, I. (2013b). Exploring the bovine rumen bacterial community from birth to adulthood. The ISME Journal, 7(6), 1069–1079.

16. Hayes, B. J., Donoghue, K. A., Reich, C. M., Mason, B. A., Bird-Gardiner, T., Herd, R. M., & Arthur, P. F. (2016). Genomic heritabilities and genomic estimated breeding values for methane traits in Angus cattle. Journal of Animal Science, 94(3), 902–908.

17. Roehe, R., Dewhurst, R. J., Duthie, C. A., Rooke, J. A., McKain, N., Ross, D. W., … & Wallace, R. J. (2016). Bovine host genetic variation influences rumen microbial methane production with best selection criterion for low methane emitting and efficiently feed converting hosts based on metagenomic gene abundance. PLoS genetics, 12(2), e1005846.

18. Rooke, J. A., Wallace, R. J., Duthie, C. A., McKain, N., de Souza, S. M., Hyslop, J. J., … & Roehe, R. (2014). Hydrogen and methane emissions from beef cattle and their rumen microbial community vary with diet, time after feeding and genotype. British Journal of Nutrition, 112(3), 398–407.

19. Martiny, J. B., Martiny, A. C., Brodie, E., Chase, A. B., Rodríguez-Verdugo, A., Treseder, K. K., & Allison, S. D. (2023). Investigating the eco-evolutionary response of microbiomes to environmental change. Ecology Letters.

20. Henderson, G., Cox, F., Ganesh, S., Jonker, A., Young, W., Global Rumen Census Collaborators, Abecia, L., Angarita, E., Aravena, P., Nora Arenas, G., Ariza, C., Attwood, G. T., Mauricio Avila, J., Avila-Stagno, J., Bannink, A., Barahona, R., Batistotti, M., Bertelsen, M. F., Brown-Kav, A., … Janssen, P. H. (2015). Rumen microbial community composition varies with diet and host, but a core microbiome is found across a wide geographical range. Scientific Reports, 5(1), 14567.

21. Petri, R. M., Schwaiger, T., Penner, G. B., Beauchemin, K. A., Forster, R. J., mckinnon, J. J., & mcallister, T. A. (2013). Characterization of the Core Rumen Microbiome in Cattle during Transition from Forage to Concentrate as Well as during and after an Acidotic Challenge. Plos ONE, 8(12), e83424.

22. Wallace, R. J., Rooke, J. A., mckain, N., Duthie, C.-A., Hyslop, J. J., Ross, D. W., Waterhouse, A., Watson, M., & Roehe, R. (2015). The rumen microbial metagenome associated with high methane production in cattle. BMC Genomics, 16(1), 839.

23. Chiri, E., Nauer, P. A., Lappan, R., Jirapanjawat, T., Waite, D. W., Handley, K. M., Hugenholtz, P., Cook, P. L. M., Arndt, S. K., & Greening, C. (2021). Termite gas emissions select for hydrogenotrophic microbial communities in termite mounds. Proceedings of the National Academy of Sciences, 118(30), e2102625118.

24. Denman, S. E., Martinez Fernandez, G., Shinkai, T., Mitsumori, M., & McSweeney, C. S. (2015). Metagenomic analysis of the rumen microbial community following inhibition of methane formation by a halogenated methane analog. Frontiers in Microbiology, 6.

25. Getahun, M. N., Villinger, J., Bargul, J. L., Muema, J. M., Orone, A., Ngiela, J., … & Masiga, D. K. (2022). Molecular characterization of pathogenic African trypanosomes in biting flies and camels in surra-endemic areas outside the tsetse fly belt in Kenya. International Journal of Tropical Insect Science, 42(6), 3729–3745.

26. Dagar, S. S., Kumar, S., Griffith, G. W., Edwards, J. E., Callaghan, T. M., Singh, R., … & Puniya, A. K. (2015). A new anaerobic fungus (Oontomyces anksri gen. nov., sp. nov.) from the digestive tract of the Indian camel (Camelus dromedarius). Fungal Biology, 119(8), 731–737.

27. Brooke, C. G., Najafi, N., Dykier, K. C., & Hess, M. (2019). Prevotella copri, a potential indicator for high feed efficiency in western steers. Animal Science Journal, 90(5), 696–701.

28. Xue, M. Y., Xie, Y. Y., Zhong, Y., Ma, X. J., Sun, H. Z., & Liu, J. X. (2022). Integrated meta-omics reveals new ruminal microbial features associated with feed efficiency in dairy cattle. Microbiome, 10(1), 32.

29. Doelman, J., mcknight, L. L., Carson, M., Nichols, K., Waterman, D. F., & Metcalf, J. A. (2019). Postruminal infusion of calcium gluconate increases milk fat production and alters fecal volatile fatty acid profile in lactating dairy cows. Journal of Dairy Science, 102(2), 1274–1280.

30. Dagar, S. S., Kumar, S., Mudgil, P., & Puniya, A. K. (2018). Comparative evaluation of lignocellulolytic activities of filamentous cultures of monocentric and polycentric anaerobic fungi. Anaerobe, 50, 76–79.

31. Saye, L. M. G., Navaratna, T. A., Chong, J. P. J., O’Malley, M. A., Theodorou, M. K., & Reilly, M. (2021). The Anaerobic Fungi: Challenges and Opportunities for Industrial Lignocellulosic Biofuel Production. Microorganisms, 9(4), 694.

32. Aristizabal-Henao, J. J., Lemas, D. J., Griffin, E. K., Costa, K. A., Camacho, C., & Bowden, J. A. (2021). Metabolomic Profiling of Biological Reference Materials using a Multiplatform High-Resolution Mass Spectrometric Approach. Journal of the American Society for Mass Spectrometry, 32(9), 2481–2489.

33. Dagaew, G., Wongtangtintharn, S., Suntara, C., Prachumchai, R., Wanapat, M., & Cherdthong, A. (2022). Feed utilization efficiency and ruminal metabolites in beef cattle fed with cassava pulp fermented yeast waste replacement soybean meal. Scientific Reports, 12(1), 16090.

34. Cho, I., & Blaser, M. J. (2012). The human microbiome: at the interface of health and disease. Nature Reviews Genetics, 13(4), 260–270.

35. Lloyd-Price, J., Abu-Ali, G., & Huttenhower, C. (2016). The healthy human microbiome. Genome medicine, 8(1), 1–11.

36. Getahun, M. N., Ngiela, J., Makwatta, J. O., Ahuya, P., Simon, T. K., Kamau, S. K., … & Masiga, D. (2022). Metabolites from trypanosome-infected cattle as sensitive biomarkers for animal trypanosomosis. Frontiers in Microbiology, 2517.

37. Cardoso-Gutierrez, E., Aranda-Aguirre, E., Robles-Jimenez, L. E., Castelán-Ortega, O. A., Chay-Canul, A. J., Foggi, G., Angeles-Hernandez, J. C., Vargas-Bello-Pérez, E., & González-Ronquillo, M. (2021). Effect of tannins from tropical plants on methane production from ruminants: A systematic review. Veterinary and Animal Science, 14, 100214.

38. Gemeda, B. S., & Hassen, A. (2014). Effect of Tannin and Species Variation on In vitro Digestibility, Gas, and Methane Production of Tropical Browse Plants. Asian-Australasian Journal of Animal Sciences, 28(2), 188–199.

39. Mohammed, A. S., Animut, G., Urge, M., & Assefa, G. (2020). Grazing behavior, dietary value and performance of sheep, goats, cattle and camels co-grazing range with mixed species of grazing and browsing plants. Veterinary and Animal Science, 10, 100154.

40. Degen, A. A., Benjamin, R. W., Mishorr, T., Kam, M., Becker, K., Makkar, H. P. S., & Schwartz, H. J. (2000). Acacia saligna as a supplementary feed for grazing desert sheep and goats. The Journal of Agricultural Science, 135(1), 77–84.

41. Kewan, K. Z., Elkhouly, A. A., Negm, A. M., & Javadi, A. (2019). Feedstock values of some common fodder halophytes in the Egyptian desert. 9.

42. El-Waziry, A. M., Basmaeil, S. M., Al-Owaimer, A. N., Metwally, H. M., Ali, M. H., & Al-Harbi, M. S. (2019). Effect of replacing alfalfa hay with acacia foliage on the growth performance, in vitro gas production and rumen fermentation in goats. Adv. Anim. Vet. Sci, 7(9), 738–744.

43. Belanche, A., Kingston-Smith, A. H., Griffith, G. W., & Newbold, C. J. (2019). A Multi-Kingdom Study Reveals the Plasticity of the Rumen Microbiota in Response to a Shift From Non-grazing to Grazing Diets in Sheep. Frontiers in Microbiology, 10, 122.

44. Van Soest, P. J., Robertson, J. B., Hall, M. B., & Barry, M. C. (2020). Klason lignin is a nutritionally heterogeneous fraction unsuitable for the prediction of forage neutral-detergent fibre digestibility in ruminants. British Journal of Nutrition, 124(7), 693–700.

45. Owens, F. N., & Basalan, M. (2016). Ruminal Fermentation. In D. D. Millen, M. De Beni Arrigoni, & R. D. Lauritano Pacheco (Eds.), Rumenology (pp. 63–102). Springer International Publishing.

46. Mansourian, S., Corcoran, J., Enjin, A., Löfstedt, C., Dacke, M., & Stensmyr, M. C. (2016). Fecal-Derived Phenol Induces Egg-Laying Aversion in Drosophila. Current Biology, 26(20), 2762–2769.

47. Ferreira, L. L., Sarria, A. L. F., de Oliveira Filho, J. G., de Silva, F. De O., Powers, S. J., Caulfield, J. C., Pickett, J. A., Birkett, M. A., & Borges, L. M. F. (2019). Identification of a non-host semiochemical from tick-resistant donkeys (Equus asinus) against Amblyomma sculptum ticks. Ticks and Tick-Borne Diseases, 10(3), 621–627.

48. Getahun, M. N., Ahuya, P., Ngiela, J., Orone, A., Masiga, D., & Torto, B. (2020). Shared volatile organic compounds between camel metabolic products elicits strong Stomoxys calcitrans attraction. Scientific reports, 10(1), 21454.

49. Spor, A., Koren, O., & Ley, R. (2011). Unravelling the effects of the environment and host genotype on the gut microbiome. Nature Reviews Microbiology, 9(4), 279–290.

50. Zhang, Q., Difford, G., Sahana, G., Løvendahl, P., Lassen, J., Lund, M. S., … & Janss, L. (2020). Bayesian modeling reveals host genetics associated with rumen microbiota jointly influence methane emission in dairy cows. The ISME journal, 14(8), 2019–2033

51. Gruninger, R. J., Puniya, A. K., Callaghan, T. M., Edwards, J. E., Youssef, N., Dagar, S. S., Fliegerova, K., Griffith, G. W., Forster, R., Tsang, A., mcallister, T., & Elshahed, M. S. (2014). Anaerobic fungi (phylum Neocallimastigomycota): Advances in understanding their taxonomy, life cycle, ecology, role and biotechnological potential. FEMS Microbiology Ecology, 90(1), 1–17.

52. Cunha, C. S., Veloso, C. M., Marcondes, M. I., Mantovani, H. C., Tomich, T. R., Pereira, L. G. R., Ferreira, M. F. L., Dill-McFarland, K. A., & Suen, G. (2017). Assessing the impact of rumen microbial communities on methane emissions and production traits in Holstein cows in a tropical climate. Systematic and Applied Microbiology, 40(8), 492– 499.

53. Foroutan, A., Fitzsimmons, C., Mandal, R., Piri-Moghadam, H., Zheng, J., Guo, A., Li, C., Guan, L. L., & Wishart, D. S. (2020). The Bovine Metabolome. Metabolites, 10(6), 233.

54. Zhu, C., Li, C., Wang, Y., & Laghi, L. (2019). Characterization of Yak Common Biofluids Metabolome by Means of Proton Nuclear Magnetic Resonance Spectroscopy. Metabolites, 9(3), 41.

55. Newbold, C. J., & Ramos-Morales, E. (2020). Ruminal microbiome and microbial metabolome: effects of diet and ruminant host. Animal, 14(S1), s78–s86.

56. Wallace, R. J., Snelling, T. J., McCartney, C. A., Tapio, I., & Strozzi, F. (2017). Application of meta-omics techniques to understand greenhouse gas emissions originating from ruminal metabolism. Genetics Selection Evolution, 49(1), 9.

57. Grossi, G., Goglio, P., Vitali, A., & Williams, A. G. (2019). Livestock and climate change: impact of livestock on climate and mitigation strategies. Animal Frontiers, 9(1), 69–76.

58. Khan, N., Ali, S., Zandi, P., Mehmood, A., Ullah, S., Ikram, M., Ismail, I., Shahid, M. A., & Babar, M. A. (2020). Role of sugars, amino acids and organic acids in improving plant abiotic stress tolerance. Pakistan Journal of Botany, 52(2).

59. Mokaya, H. O., Nkoba, K., Ndunda, R. M., & Vereecken, N. J. (2022). Characterization of honeys produced by sympatric species of Afrotropical stingless bees (Hymenoptera, Meliponini). Food Chemistry, 366, 130597.

60. Clauss, M., Kaiser, T., & Hummel, J. (2008). The Morphophysiological Adaptations of Browsing and Grazing Mammals. In I. J. Gordon & H. H. T. Prins (Eds.), The Ecology of Browsing and Grazing (Vol. 195, pp. 47–88). Springer Berlin Heidelberg.

61. Malheiros, J. M., Correia, B. S. B., Ceribeli, C., Cardoso, D. R., Colnago, L. A., Junior, S. B., Reecy, J. M., Mourão, G. B., Coutinho, L. L., Palhares, J. C. P., Berndt, A., & de Almeida Regitano, L. C. (2021). Comparative untargeted metabolome analysis of ruminal fluid and feces of Nelore steers (Bos indicus). Scientific Reports, 11(1), 12752.

62. Ward, D., Schmitt, M. H., & Shrader, A. M. (2020). Are there phylogenetic differences in salivary tannin-binding proteins between browsers and grazers, and ruminants and hindgut fermenters? Ecology and Evolution, 10(19), 10426–10439.

63. Poulopoulou, I., & Hadjigeorgiou, I. (2021). Evaluation of Terpenes’ Degradation Rates by Rumen Fluid of Adapted and Non-adapted Animals. Natural Products and Bioprospecting, 11(3), 307–313.

64. Tarasov, D., Leitch, M., & Fatehi, P. (2018). Lignin–carbohydrate complexes: Properties, applications, analyses, and methods of extraction: a review. Biotechnology for Biofuels, 11(1), 269.

65. Urrutia, N. L., & Harvatine, K. J. (2017). Acetate Dose-Dependently Stimulates Milk Fat Synthesis in Lactating Dairy Cows. The Journal of Nutrition, 147(5), 763–769.

66. De Paula, E. M., Samensari, R. B., Machado, E., Pereira, L. M., Maia, F. J., Yoshimura, E. H., Franzolin, R., Faciola, A. P., & Zeoula, L. M. (2016). Effects of phenolic compounds on ruminal protozoa population, ruminal fermentation, and digestion in water buffaloes. Livestock Science, 185, 136–141.

67. Mahfuz, S., Shang, Q., & Piao, X. (2021). Phenolic compounds as natural feed additives in poultry and swine diets: A review. Journal of Animal Science and Biotechnology, 12(1), 48.

68. Rossi, R., Stella, S., Ratti, S., Maghin, F., Tirloni, E., & Corino, C. (2017). Effects of antioxidant mixtures in the diet of finishing pigs on the oxidative status and shelf life of longissimus dorsi muscle packaged under modified atmosphere1, 2. Journal of Animal Science, 95(11), 4986–4997.

69. Machado, G., Santos, F., Lourega, R., Mattia, J., Faria, D., Eichler, P., & Auler, A. (2020). Biopolymers from lignocellulosic biomass: feedstocks, production processes, and applications. Lignocellulosic biorefining technologies, 125–158.

70. Guliński, P. (2021). Ketone bodies – causes and effects of their increased presence in cows’ body fluids: A review. Veterinary World, 1492–1503.

71. Tawich, S. K., Bargul, J. L., Masiga, D., & Getahun, M. N. (2021). Supplementing Blood Diet with Plant Nectar Enhances Egg Fertility in Stomoxys calcitrans. Frontiers in Physiology, 12, 646367.

72. Tamura, K., Stecher, G., & Kumar, S. (2021). MEGA11: Molecular Evolutionary Genetics Analysis Version 11. Molecular Biology and Evolution, 38(7), 3022–3027.

73. Omondi, V. O., Bosire, G. O., Onyari, J. M., & Getahun, M. N. (2022). A Comparative Investigation of Volatile Organic Compounds of Cattle Rumen Metabolites using HS-SPME and porapak-Q Odor Trapping Methods. Analytical Chemistry Letters, 12(4), 451– 459.

74. R Core Team (2022). R: A language and environment for statistical computing. R Foundation for Statistical Computing, Vienna, Austria. URL http://www.R-project.org/.

75. Hammer, O., Harper, D. A. T., & Ryan, P. D. (2001). PAST: Paleontological Statistics Software Package for Education and Data Analysis.

76. Martin, M. (2011). Cutadapt removes adapter sequences from high-throughput sequencing reads. EMBnet. Journal, 17(1), 10–12.

